# Poly(A)-ClickSeq: click-chemistry for next-generation 3´-end sequencing without RNA enrichment or fragmentation

**DOI:** 10.1101/109272

**Authors:** Andrew Routh, Ping Ji, Elizabeth Jaworski, Zheng Xia, Wei Li, Eric J. Wagner

## Abstract

The recent emergence of alternative polyadenylation (APA) as an engine driving transcriptomic diversity has stimulated the development of sequencing methodologies designed to assess genome-wide polyadenylation events. The goal of these approaches is to enrich, partition, capture, and ultimately sequence poly(A) site junctions. However, these methods often require poly(A) enrichment, 3´ linker ligation steps, and RNA fragmentation, which can necessitate higher levels of starting RNA, increase experimental error, and potentially introduce bias. We recently reported a click-chemistry based method for generating RNAseq libraries called “ClickSeq”. Here, we adapt this method to direct the cDNA synthesis specifically toward the 3´ UTR/poly(A) tail junction of cellular RNA. With this novel approach, we demonstrate sensitive and specific enrichment for poly(A) site junctions without the need for complex sample preparation, fragmentation or purification. Poly(A)-ClickSeq (PAC-seq) is therefore a simple procedure that generates high-quality RNA-seq poly(A) libraries. As a proof-of-principle, we utilized PAC-seq to explore the poly(A) landscape of both human and *Drosophila* cells in culture and observed outstanding overlap with existing poly(A) databases and also identified previously unannotated poly(A) sites. Moreover, we utilize PAC-seq to quantify and analyze APA events regulated by CFIm25 illustrating how this technology can be harnessed to identify alternatively polyadenylated RNA.

## Introduction

With the exception of replication-dependent histone mRNA, poly(A) tails are ubiquitous to all eukaryotic mRNAs and function to stimulate translation and impart protection from cellular exonucleases (reviewed in (1)). Not surprisingly, the 3´ termini of many RNA viruses, including picornaviruses (2) and HIV(3), have also been found to possess poly(A) tails. Cellular mRNA receive poly(A) tails through the process of cleavage and polyadenylation where the pre-mRNA is co-transcriptionally cleaved and subsequently used as a substrate for poly(A) polymerase. The location of cleavage and polyadenylation near the 3´ end of a pre-mRNA is governed by three primary sequence elements: the hexameric polyadenylation signal (PAS, typically AWUAAA)(4), the cleavage site (typically a CA dinucleotide), and the downstream sequence element (DSE, typically U/UG rich). The collective adherence to consensus that these three elements possess is thought to dictate the overall efficiency of cleavage and polyadenylation at a particular site (5). The enzymatic process of cleavage and polyadenylation is carried out by a group of proteins called the cleavage and polyadenylation (CPA) complex that contains at least fifteen subunits, the core members of which are conserved from yeast to humans (reviewed in (6)). Complete loss of activity of any of these core CPA subunits leads to broad failure to produce mRNA ultimately resulting in loss of cell viability.

While initially thought to be a constitutive or house-keeping event, recent work from many laboratories have shown that cleavage and polyadenylation is highly dynamic (reviewed in (7)). Underscoring its importance, it has been observed that greater than 50% of mammalian mRNA have multiple potential cleavage and polyadenylation sites giving rise distinct mRNA isoforms of different length(8). This process, termed alternative polyadenylation (APA) dramatically increases the known diversity of the eukaryotic transcriptome (reviewed in (9,10)). The preponderance of data demonstrates that APA is developmentally regulated(11,12), can occur as tissues become more differentiated(13,14), when they are subject to cellular stress(15), or during diseased states such as cellular transformation(16). In particular it has been shown that when cells are induced to proliferate and/or undergo cellular transformation, there is a global trend toward the selective use of proximal poly(A) signals (pPAS) resulting in the production of mRNA with truncated 3’UTR that are not effectively targeted by miRNA(17,18). The mechanisms that manage APA regulation are less clear and several factors have been identified that can influence poly(A) site selection including chromatin or DNA modification(19,20), changes in RNA polymerase II elongation efficiency(21), and modulation of RNA binding/processing factors that are known to play a role in cleavage and polyadenylation(22-29). Of the CPA machinery, either increases in CstF64 expression (30) or decreases in CFIm complex member levels (23,24,31) leads to broad shortening of 3´UTRs suggesting that these two factors may play antagonistic roles in governing poly(A) site selection.

In light of the recent appreciation for APA, profiling the position of the poly(A) tail using high-throughput sequencing technologies is critical to understand the complex interplay of poly(A) tail location with mRNA stability, degradation and translation. In the simplest manner, the positions of poly(A) tails can be directly extracted from both short-read RNA-seq and long-read nanopore or Pacbio (e.g. IsoSeq) sequencing by extracting non-reference ‘A’s from mapped sequence reads(32). Alternatively, approaches have been developed that infer poly(A) tail position and abundances through computational analysis of standard RNA-seq using designer algorithms catered to measure the relative density of sequence reads within the 3´UTR relative to that observed in the coding regions(33). The advantage of these approaches is that they only require standard RNA-seq analysis and can be employed retrospectively onto existing datasets. However, they have the disadvantage in that precise poly(A) site junctions are not enriched relative to the rest of the transcriptomic data and so datasets are invariably large and require high depth sequencing runs (>100M reads) as only a subset of the RNA-seq will contribute to the analysis.

As a result, a number of strategies have been developed with the specific goal of enriching for the junction of the encoded 3´UTR ends and the beginning of the non-templated poly(A) tail (11,13,34–41). Common themes found in several of these techniques are the enrichment for poly(A)+ RNA from total RNA, fragmentation of mRNA using a variety of approaches (e.g. enzymatic, heat, sonication), and attachment of an adaptor to the 3´ end either through the use of a splinted oligo or directly to the terminus of the poly(A) tail. These initial steps can also involve the use of a biotin-containing oligonucleotide to allow for purification of the desired library intermediates using streptavidin magnetic beads. These approaches typically utilize between 1–20M reads and have the advantage of allowing precise mapping of the position of the poly(A) tail addition. However, these approaches often entail complex experimental pipelines and purification strategies that can impart sample bias and reduce throughput capacity. Importantly, these challenges can reduce the number of core facilities offering these types of sequencing technologies thereby limiting their application only to laboratories with more than routine experience in sequencing library preparation.

Here we present a novel approach that provides a number of advantages over other methodologies due to its simplicity, cost-effectiveness, and speed while providing high-quality, unbiased sequencing libraries. Our approach is a subtle alteration of an RNA-seq technique we recently reported called ‘ClickSeq’(42). For poly(A)-ClickSeq (PAC-seq), small amounts of three 3´-azido-nucleotides (AzGTP, AzCTP and AzATP) are added to oligo-dT primed cDNA synthesis reactions yielding cDNA fragments that are stochastically terminated upstream of the 3´UTR/poly(A) junction, but not within the poly(A) tail. Subsequently, the azido-terminated cDNA can be purified, ‘click-ligated’ to an alkyne-functionalized 5’ Illumina adaptor and an NGS library enriched with 3´UTR/poly(A) junctions is then created by standard PCR. As a demonstration of its applicability, we use PAC-Seq to analyze total cellular RNA from HeLa cell extracts and demonstrate that this approach is robust and can thoroughly capture authentic pre-validated polyadenylated sites without the need for any sample purification, enrichment or fragmentation. Moreover, this can be achieved with a minimal number of extraneous sequence reads allowing for experiments with multiple replicates to be loaded into a single flowcell of an Illumina MiSeq. We also analyzed multiple replicates of HeLa cells that have been depleted of CFIm25 to demonstrate the ability of PAC-seq to identify and quantify APA regulation. Finally, we characterize the poly(A) site profile of *Drosophila* S2 cells in culture and found that depletion of fly orthologue of CFIm25 (CG3689) induces only a small number of APA changes, indicating that the role of CFIm25 in regulated *Drosophila* APA may not be as extensive in fly. Overall, the simplicity, cost-effectiveness and fast turnaround of PAC-Seq will allow investigation into a wide-range of complex samples that were previously either too uneconomical or intractable to analyze.

### MATERIALS AND METHODS

#### Isolation of RNA from HeLa cells with si-RNAs for CFIm25

Parental HeLa cells were purchased from ATCC (Cat#CCL-2) and maintained in Eagle’s Minimum Essential Medium (Lonza, Cat#12-604F) with 10% fetal bovine serum. The cells are transfected with three different siRNAs for CFIm25 (Sigma Aldrich, St. Louis, MO, ID: SASI_Hs01_00146875~77) and negative control siRNA (Sigma Aldrich, St. Louis, MO, ID:SIC002) using previously established approaches (43). Knockdown of CFIm25 was determined by Western blotting with anti-CFIm25 antibody (Proteintechlab, Rosemont, IL, Cat# 10322-1-AP), GAPDH (Sigma, St. Louis, MO, G9545) served as a loading control. Total RNA was extracted using TRIzol Reagent (Life Technologies) using the manufacturers protocol.

#### Isolation of RNA from S2 cells with dsRNA for CFIm25

*Drosophila* S2 cells were cultured in Schneider’s Drosophila media (GIBCO) supplemented with 10% FBS, 50 units/ml penicillin, and 50 μg/ml streptomycin at 28°C. To knockdown CFIm25 in S2 cells, an individual DNA fragment in exon 1 of CFIm25 308bp in length was PCR amplified. Each primer used in the PCR contained a 5´ T7 RNA polymerase binding site (GAATTAATACGACTCACTATAGGG) followed by sequences specific for CFIm25 gene (Forward primer:+AGCGCTGGACAGAAAAGTGT and reverse primer: +CGCCTGGTTGGTGTACTTCT). The PCR products were purified and used as templates to produce dsRNA using T7 RNA polymerase (Ambion). The dsRNA products were ethanol-precipitated and resuspended in water. The dsRNAs were annealed by incubation at 65°C for 30 min followed by slow cooling to room temperature. S2 cells were incubated with dsRNA for CFIm25 or negative control dsRNA for LacZ or for three days with three hits. Total RNA was extracted using TRIzol Reagent (Life Technologies) using the manufacturers protocol. For quantitative Real Time-PCR (qRT-PCR) the mRNA was reverse transcribed using MMLV-RT (Invitrogen) using the manufacturer’s protocol to generate cDNA. The qRT-PCR reactions were performed using Stratagene MxPro3000P (Agilent Technologies) and SYBRGREEN (Fermentas). The forward primer AGGGCCTCAAGAGATTGCTA is in exon2 boundary of CFIm25 and the reverse primer ATCGTGTCCTCAACAATCCA is located in exon 3 of CFIm25. The *Drosophila* housekeeping gene ribosomal protein S17 (Rps17) served as an internal control.

#### Library Preparation

No additional purification or selection of total RNA is required as the RT primer selects for polyadenylated RNAs. Up to 4ug of total RNA was used to generate the Poly(A)-ClickSeq libraries. Reverse transcription was performed using standard protocols with the addition of spiked-in azido-nucleotides (AzVTPs). Specifically, a 1:5 5mM AzVTP:dNTP working solution was made by adding 10 μL of 10mM dNTPs to 2 μL each of 10mM AzATP, AzCTP, and AzGTP (no AzTTP) and water to a final volume of 20 μL. To begin, 4ug RNA, 1 μL of 5mM AzVTP:dNTPs working solution, and 1 μL 50μM 3´Illumina_4N_21T primer (GT GACTGGAGTT CAGACGT GT GCT CTTCCGAT CTNNN NTTTTTTTT TTTTTTTTTTTTT) were mixed in 13 μL total volume and was heated to 95°C for 2 min to denature the RNA then snap cooled on ice, >1min. (NB: This is a non-anchored poly-T primer.) Superscript III Reverse Transcriptase (Invitrogen), 5X Superscript First Strand Buffer, DTT, and RNase OUT (Invitrogen) was added for 20 μL total final volume and the reaction was incubated at 50° for 20 min, then 75° for 15 min. Room temperature incubation was avoided during mixing of components to avoid non-specific amplification. After cDNA synthesis, the template RNA was removed with the addition of 10U RNase H (NEB) incubated at 37° for 20 mins. Next, the azido-terminated cDNA was purified using the Zymo DNA Clean and Concentrator Kit (Cat #11-303C) and eluted with 10 μL of 50 mM HEPES pH 7.2.

#### Click-Reaction

The ‘Click-Adapter’ (5’ Hexynyl-NNNNAGATCGGAAGAGCGTCGTGTAGGGAAAGAGTGTAGATCTCGGTGGTCGCCGTATCATT) was added onto the azido-terminated cDNA by copper-catalyzed alkyne-azide cycloaddition (CuAAC) (42). The click-reaction was made by diluting all 10 μL of the azido-terminated cDNA in 20μL 100% DMSO, 3 μL 5 μM Click-Adapter and catalyzing the reaction twice with 0.4 μL 50mM Vitamin C and 2 μL 10mM Cu-TBTA (Lumiprobe) for 30 min at room temperature. The clicked-linked cDNA was then purified on a Zymo DNA column.

#### PCR Amplification

The final PCR amplification appends the remaining Illumina adapters and the desired demultiplexing index. Reactions were set up with the following reaction components: 5 μL Click-ligated cDNA, 2.5 μL 5 μM Indexing primer (CAAGCAGAAGACGGCATACGAGATnnnnnnGTGACTGGAGTTCAGACGTGT, where nnnnnn is the sequence of the desired index), 2.5 μL 5 μM Short Universal Primer (AATGATACGGCGACCACCGAG), and 25 μL 2X One Taq Standard Buffer Master Mix for a final 50 μL reaction. Optimized thermocycler conditions are as follows: 94° 4 min; 53° 30 sec; 68° 10 min; [94° 30 sec, 53° 30 sec, 68° 2 min] × 20−22; 68° 5 min. Amplified PCR product was then run on a 2% precast agarose e-gel (Invitrogen, E-Gel Electrophoresis System) for 10 minutes and ~200-300bp fragments (for 1×150 SE Illumina) or ~200-400bp fragments (for 1×250 SE Illumina) were excised and cleaned using the Zymo Research Gel DNA Recovery Kit. Final yield of size selected cDNA library was quantified using a QuBit fluorimeter.

#### Sequencing

Libraries were pooled and sequenced using the manufacturer’s standard operating procedures on either a HiSeq 1500 using a HiSeq Rapid SBS kit v2 obtaining 1×250bp SE reads, or a MiSeq using a MiSeq Reagent Kit v2 (300 cycles) obtaining 1×250bp SE reads. Raw data was demultiplexed using TruSeq indexes using the CASAVA pipeline or MiSeq Reporter Software. All read data can be accessed through the GEO database (GSE94950).

#### Read Processing and Quality Filtering

All custom python scripts (as well as example batch recipes and instructions) used in the following read-processing steps are available **in Supplementary Datafile 1**. Raw reads were trimmed to remove TruSeq adaptors and the first 6 nucleotides derived from the ‘Click - Adaptor’ adaptors using *cutadapt*(*44*); variables: -a nnnnagatcggaagagc -m 60. We discarded reads shorter than 60 nucleotides as these would be too short to yield both a poly(A) tail as well as sufficient nucleotides to provide an unambiguous mapping. Next, *cutadapt* was used a second time to search for reads containing poly(A) tails at least 15nts in length, allowing for one mismatch; variables: -b AAAAAAAAAAAAAAA -n 2 -O 6 -m 40. Using a custom script (**Supplementary Datafile 1**), the poly(A) tail length is extracted by comparing the de-adenylated reads to the pre-trimmed reads and this information is appended to the read name of the data file. The trimmed, de-adenylated reads were additionally quality filtered using the *fastxtoolkit* (http://hannonlab.cshl.edu/fastxtoolkit/) to ensure that >98% of the nucleotides in each read had a PHRED score greater than 20. This process yields single-end reads without poly(A)s at least 40 nts in length.

#### Read Mapping and Poly(A) site annotation

The processed reads were mapped using the *Hisat2*(45) splice-aware aligner to the reference human genome (hg19) or *Drosophila melanogaster* (dm6) using the default mapping parameters, with the exception of disallowing soft-pads at the 3´ end of the mapped read in order to prevent mis-annotation of the poly(A)site; variables: --sp 3,7. The position of the poly(A) tails are given by the final nucleotide of the mapped reads. This locus, the number mapped reads and the number of A’s present in each mapped read are written to BEDGraph files using custom scripts (**Supplementary Data 1**). The BEDGraph contains an extra non-canonical entry comprising an data array whose coordinate (1-300) corresponds to poly(A) length and the value at that coordinate returns the number of reads that had that poly(A) length. This information allows us to apply a filter requiring each unique poly(A) tail to contain non-primer/non-templated A’s as well as multiple mapping reads.

A range of values for this filter were tested requiring between 1 and 50 reads per event and requiring an average of between 1 and 10 non-templated A’s (22 to 31 total As). The number of reads retained after this filter is illustrated in the heat map in **Supplemental Figure 1**. The number of retained events drops quickly as a function of the number reads required plateauing at approximately N=5, and modestly as a function of poly(A) length. Therefore, we filtered these sites requiring each site to have at least 5 mapped reads and the poly(A) tails to have at least 5 non-templated As per reads. Finally, the base composition of 20 nts nucleotides from the reference genome downstream of the poly(A) sites were inspected using *samtools* (46) and custom scripts (**Supplementary Data 1**). The frequency of nucleotides found in this regions are shown in Supplementary Figure 2. This revealed an abundance of sites that were predominantly As downstream of the poly(A) sites in the reference genome, and are therefore likely to be internally primed sites, rather than bona fide poly(A) tails. Additionally, T’s were also found to be over-represent in these regions, consistent with the observation that U-rich tracts promote pre-mRNA cleavage (5). Therefore, only Poly(A) sites containing 15 or fewer A’s in these sequences were used for further analysis. BEDgraph files containing the poly(A) sites identified in this manuscript are available in **Supplementary Datafiles 2-4**.

For alternative poly-adenylation analysis, multiple poly(A) sites occurring within 10nts of one another were clustered into a single site, with the frequency of the clustered site equaling the sum of the individual sites. Sites found within the terminal exon of genes annotated in the UCSC genome browser were extracted and compared between wild-type and CF25Im knock-down cell-lines. If multiple poly(A) sites were found within the terminal exon and if the relative usage of these was altered by greater than 10% between the wild-type and knock-down cell types then these poly(A) sites were deemed to be alternatively polyadenylated.

#### Motif Enrichment Analysis

The sequences from the reference genome either upstream or downstream of the poly(A) sites were extracted using *samtools* (46) and custom scripts (**Supplementary Data 1**). Unique sequences were searched for RNA motif enrichment using the *dreme* (47) component of the MEME suite; variables: -rna –norc -mink 4 -maxk 8.Following this analysis, the distribution probability of enriched motifs were determined using *CentriMo* (*48*); variables: --norc.

### RESULTS

#### Poly(A)-ClickSeq Library Generation

We recently reported a technique called ‘ClickSeq’ that uses azido-nucleotide terminators in randomly-primed RT reactions to produce cDNA fragments from non-fragmented template RNA (42). Azido-nucleotides are stochastically incorporated during cDNA synthesis inducing chain-termination yielding a distribution of cDNA fragment lengths, which is determined by the ratio of AzNTPs to dNTPs. As a result of chain termination, the cDNA fragments are blocked by an azido-group at their 3´ end. Using copper-catalyzed azide-alkyne cycloaddition (CuAAC) (49), we demonstrated that we could ‘click-ligate’ 5´-hexynyl functionalized DNA oligos corresponding to the Illumina universal sequencing primer onto these 3´-azido-terminated fragments, generating unnatural triazole-linked ssDNA molecules. Importantly, these ssDNA templates are bio-compatible (50). Therefore, with a standard PCR reaction we can amplify these fragments to generate high-quality Illumina sequencing libraries with even sequence coverage (51). Moreover, this approach provides many advantages over many over RNA-seq methodologies due to its simplicity, the removal of the fragmentation and ligation steps, and the reduction of artifactual RNA recombination (42).

Here, we sought to utilize this approach to target sequencing to only the 3’ ends of polyadenylated RNAs: “Poly(A)-ClickSeq”; or PAC-seq. For PAC-seq, rather than using a random primer, we initiate reverse transcription using oligo(dT) primers without a non-T anchor. This primer also contains an overhang corresponding to a portion of the Illumina p7 adaptor (illustrated in **Fig. 1A**). By priming directly from poly(A) tails, we can specifically reverse transcribe polyadenylated RNAs directly from crude RNA extracts without any prior sample purification or poly(A) enrichment. Moreover, by avoiding the use of a non-anchored oligo(dT) primer, in principle the primer can anneal anywhere with the poly(A) tail. Therefore, complementary cDNA transcripts will contain ‘T’s derived from the template as well as 21 ‘T’s derived from the RT-primer. During computational read processing, this information is extracted and used to provide a quantitative assessment of mapping reliability and allow us to filter out mis-primed events that only have T’s derived from the primer and none from a poly(A) tail (**SFig. 3**).

**Figure 1:**
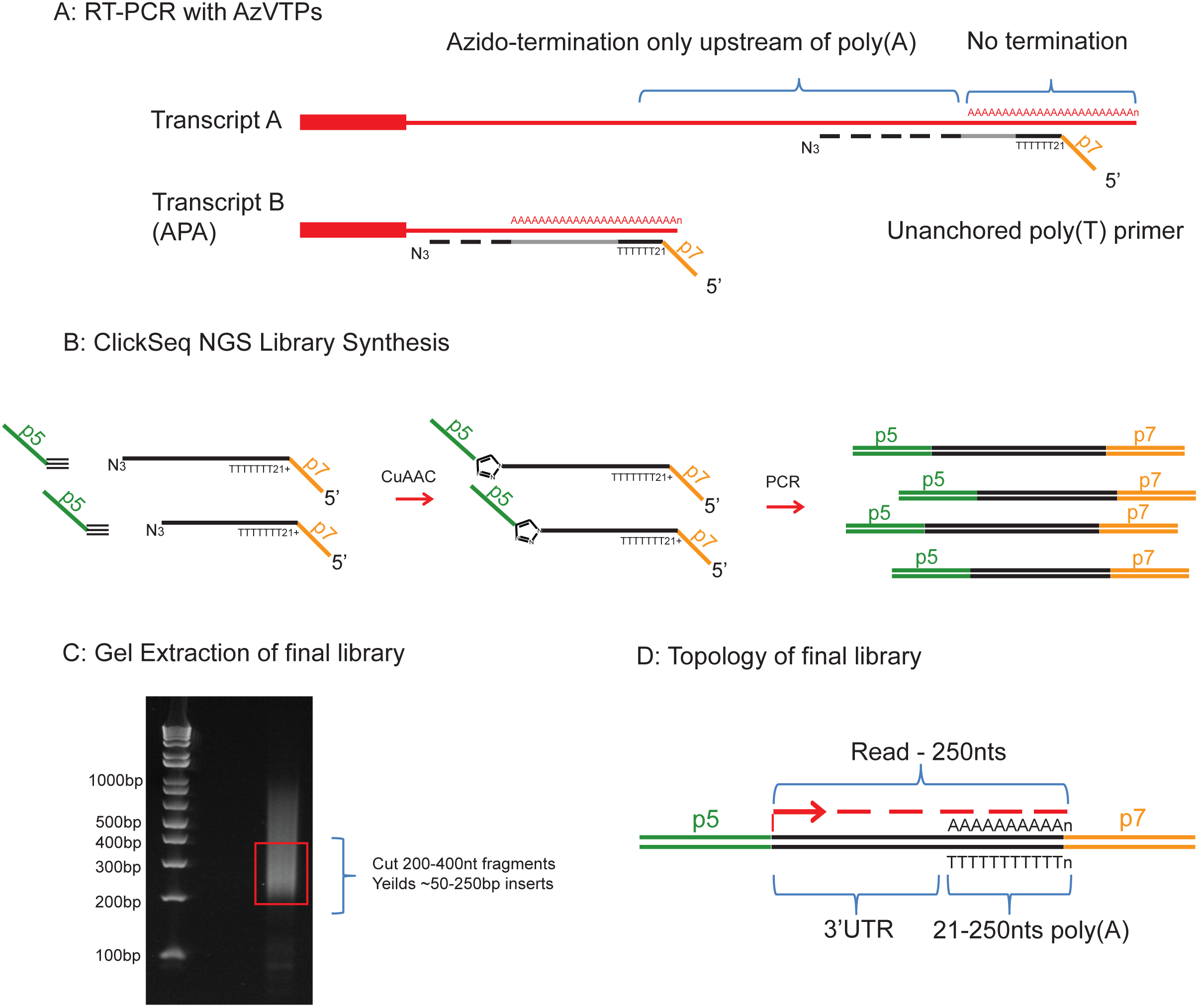
Schematic overview of Poly(A)ClickSeq pipeline. **A)** RT-PCR is launched from a nonanchored Poly(T) primer containing a portion of the Illumina p7 adaptor. RT-PCR is performed in the presence of AzATP, AzGTP and AzCTP, but not AzTTP, thus only allowing chain termination to occur upstream of the poly(A) tail in the 3’UTR. **B)** 3’-Azido-blocked cDNA fragemtns are ‘click-ligated’ to 5’hexynyl –functionalised DNA oligos containing the p5 illumine adaptor. The yields triazole-linked ssDNA whicih can be PCR-amplified using primers to the p5 and p7 Illumina adaptors. **C)** The cDNA library is analysed by gel electrophoresis and should consist of a smear of DNA products centered around 200-300bp. Appropriate cDNA fragment sizes are cut out of the gel and purified to yield a final library. D) The final lbrary consists of DNA fragments containing the illumine p5 adaptor, a portion of the 3’UTR, a stretch of As derived from both the RNA template and the poly(T) primer, and finally the p7 Illumina Indexing primer.

In ‘ClickSeq’, cDNA synthesis can terminate opposite any nucleotide. In PAC-seq, however, the critical innovation required to specifically sequence the junctions of RNA 3´UTRs and their poly(A) tails is to omit AzTTP from the reaction mixture (i.e. we provide a mixture of AzVTPs and dNTPs). Without AzTTP present in the RT-PCR reaction mixture, reverse-transcription cannot terminate opposite an ‘A’ in the RNA template. Rather, reverse-transcription must continue until non-A residues are found (**Fig. 1A**). Therefore, cDNA synthesis is stochastically terminated at a distance upstream of the 3´UTR/poly(A) junction tailored by adjusting the ratio of AzVTPs to dNTPs. This design allows for cDNA chain termination to occur only in the residues just upstream of poly(A) tail, essentially ‘homing-in’ on the junction of the 3’UTR and the poly(A) tail. We have found that a ratio of 1:5 AzVTPs:dNTPs reliably yields cDNA fragments ranging from 50-400 nts in length (42).

To finalize PAC-Seq libraries, we purify the azido-terminated cDNA, ‘click-ligate’ the 5´ Illumina adaptor, and then PCR amplify an NGS library containing the desired demultiplexing indices (**Fig. 1B**). The total size of all the adaptors including the oligo(dT) primer is 150bp. Therefore, cutting cDNA fragments 200-400nt in length will yield inserts 50-250nts in length (**Fig. 1C**). Each of the cDNA fragments will therefore contain: the full Illumina p5 adaptor; cDNA corresponding to the 3´UTR of the RNA transcript, the length of which is determined by the stochastic termination of RT-PCR; the poly(A) tail; and finally the Illumina p7 indexing adaptor (**Fig. 1D**). For optimal yield of reads containing poly(A) tails, libraries must be carefully size selected depending upon the sequencing platform and length of reads sequenced. Sequencing is initiated from the p5 adaptor. Therefore, if fragments are too large and the cDNA insert is longer than the length of the sequencing read, the 3´UTR/poly(A) tail junction will not be reached.

#### Poly(A)-ClickSeq reveals the location and relative abundance of poly(A) sites

To test our approach for the mapping of poly(A) tails, we performed 3 replicate PAC-Seq library preparations from total cellular RNA extracted from HeLa cells. HeLa cells have been well-characterized previously, both by our group and by others, and so provide a robust dataset against which to compare our mapping results. Final libraries were size-selected for fragment lengths up to 250 nts. This allows the detection of a wide range of poly(A) tail lengths. The three libraries were sequenced on a HiSeq 1500, yielding 26-36 Million raw reads per sample. These raw reads were processed as described in *Methods.* Greater than 46% of the raw demultiplexed read data were successfully processed using our pipeline, passing quality filters and containing poly(A) tails greater than 25 nts in length (**STable 1**). Therefore, our technique efficiently utilizes the data generated to find poly(A) tails. Using the splice-aware aligner, *HiSat2* (*45*), 95-97% of the processed reads from each sample were successfully mapped to the human genome (hg19) (**STable 1**). An example of the mapped PAC-Seq reads to the human gene Akt1 is shown alongside previously obtained RNA-seq coverage data of HeLa cells (31) (**Fig. 2A**). This illustrates how the PAC-Seq data is concentrated at the 3´ end of the RNA transcript. In contrast, the RNA-seq data is spread across the length of the mature mRNA.

**Figure 2:**
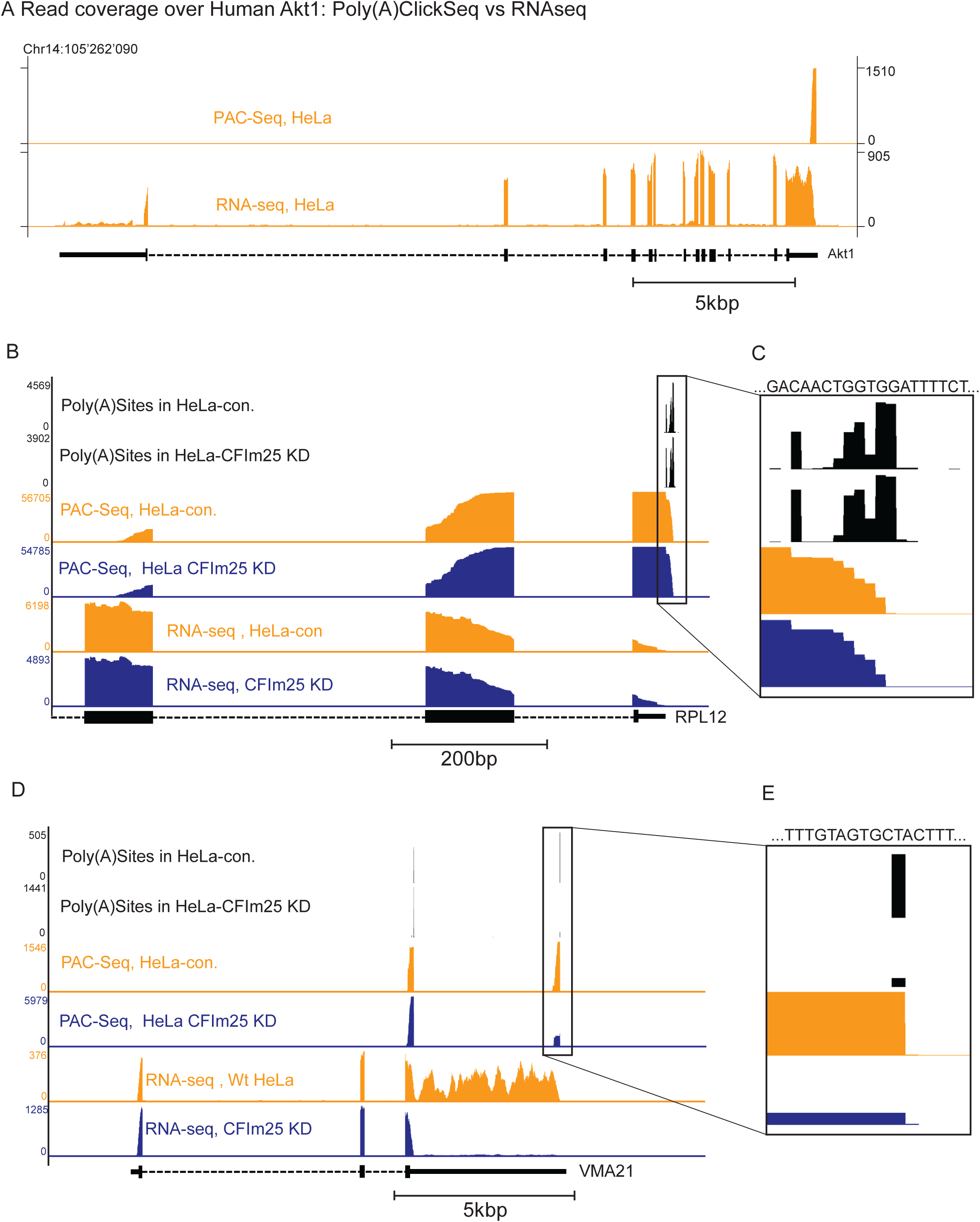
Examples of Poly(A)ClickSeq mapping data over mRNAs and comparison to RNAseq data. The positions of Poly(A) sites were determined from Poly(A)ClickSeq analysis of wild-type and CF25Im knockdown HeLa cells. UCSC tracks are shown for the Poly(A)Sites (derived from **Supplementary Datafile 2**), the coverage of the mapped Poly(A)ClickSeq reads. and from RNAseq analysis of HeLa cells from a previous analysis (REF). Scale bars are indicated below each group of tracks (NB different scale). **A)** The mapping of reads over the human gene Akt1 is illustrated. Poly(A)ClickSeq reads are only found at the very 3’ end of the mRNA transcript. In contrast, standard RNAseq coverage is distributed over all the exons. **B)** Poly(A)ClickSeq and RNAseq coverage is illustrated over the abundant transcript, RPL12 (Homo sapiens ribosomal protein L12, mRNA). PAC-Seq reveals the exact site of the poly(A) tail with nucleotide resolution revealing that the precise cleavage sites is variable. The boxed region in **B)** is enlarged in **C)** to illustrate diversity of Poly(A)Sites. **D)** Poly(A)ClickSeq and RNAseq coverage is illustrated over the alternatively poly-adenylated transcript, VMA21 (Homo sapiens VMA21 vacuolar H+-ATPase homolog (S. cerevisiae) (VMA21), mRNA). PAC-Seq reveals two poly(A) sites (the distal and proximal site). Upon CF25Im knock-down, the proximal site is significantly enriched. This observation is also supported by the RNAseq data that shows reduced read coverage over the 3’UTR of VMA21 after CF25Im KD. The boxed region in **D)** is enlarged in **E)** to illustrate distal Poly(A) Site.

From the mapped data, we can definitively determine the exact location of poly(A) tail addition. Moreover, as non-primer derived ‘A’s are found in the read data, we can also determine the distribution of poly(A) tails lengths found among the reads mapping at each specific location. With this information, we can filter the mapped reads requiring them to contain a user-defined number of ‘A’s as described in the *Methods* sections. We found that by requiring five or more reads each with five or more non-primer derived ‘A’s removed a large number of poorly-populated and likely non-specific RT-PCR products (**SFig. 1**). Interestingly, the broad distribution of poly(A) tail lengths found throughout all the mapped reads matches trends previously reported (41) (**SFig. 3**).

#### Application of Poly(A)-ClickSeq to analyze Alternative Polyadenylation

We sought to further validate the utility of PAC-seq by testing its ability to detect alternative polyadenylated sites. We and others have previously demonstrated that CFIm25 is a critical factor in the regulation poly(A) selection in mRNAs(23,28,43). Knock-down of CFIm25 results in the broad shortening of multiple mRNAs targets genome-wide. Therefore, we performed replicate CFIm25 siRNA knock-downs in HeLa cells (**SFig. 4A**), extracted total cellular RNA and prepared PAC-Seq libraries.

In total, our analysis yielded 56’937 putative poly(A) sites in the wild-type HeLa cells, and 76’176 sites in the CFIm25 KD cells (**STable 1**). By requiring sites to be found in at least two out of three replicates, we found 24’937 and 33’008 sites respectively (**STable 1**). So while specificity is greatly increased by leveraging the replicate data, the sensitivity is also decreased – resulting in the loss of over 75’000 putative poly(A) sites. Therefore the choice, implementation and interpretation of the number of replicates required in such transcriptomic analyses must be carefully considered and balanced (52). As one of the possible applications of PAC-seq is to characterize and discover any putative or novel poly(A) sites, we proceeded to analyze poly(A) sites found in two or more replicates in order to maximally utilize our data, while retaining a reasonable degree of confidence.

In the case of the highly expressed RPL12 gene that has not been found to undergo APA, we can see that the exact identity of the 3´UTR/poly(A) tail junction can vary by approximately 10 nts in either dataset (**Fig. 2B** and **2C**). This may reflect a lack of specificity or degeneracy in the selection of the 3´ cleavage site by the CPA. Nonetheless, the location of the poly(A) site found here agrees with the annotations from the UCSC database (53) as well as the anticipated poly(A) site determined from the mapping coverage in the RNA-seq data (33). VMA21 has previously been identified as subject to CFIm25 regulation and that usage of the proximal poly(A) site (pPAS) is enriched in the CFIm25 KD cells(31). Indeed, we observed similar results of reduced read density within the VMA 3´UTR upon CFIm25 KD and a clear switch from distal poly(A) site (dPAS) to pPAS usage when analyzed using PAC-seq (**Fig. 2D** and **2E**).

The majority of the detected polyadenylation events mapped to known genes in the UCSC database (~88.5%) and indeed the majority of these to annotated terminal exons as would be expected (**STable 2** and **Fig. 3A**). A further 7.53% of the detected poly(A) sites mapped within 500 nts downstream of genes in UCSC database, indicating that a substantial number of poly(A) sites in fact can be found slightly downstream of the annotated mRNA termination sites. From the remaining 992 events (3.98%) not mapping over any known annotation, 791 of these were found to have the AWUAAA motif within 100 nts upstream of the detected poly(A) sites indicating that the PAC-seq dataset is locating novel and likely *bona fide* poly(A) sites bearing canonical regulatory elements. These trends are also quite similar in the CFIm25 KD dataset (**STable 2** and **Fig. 3A**).

**Figure 3:**
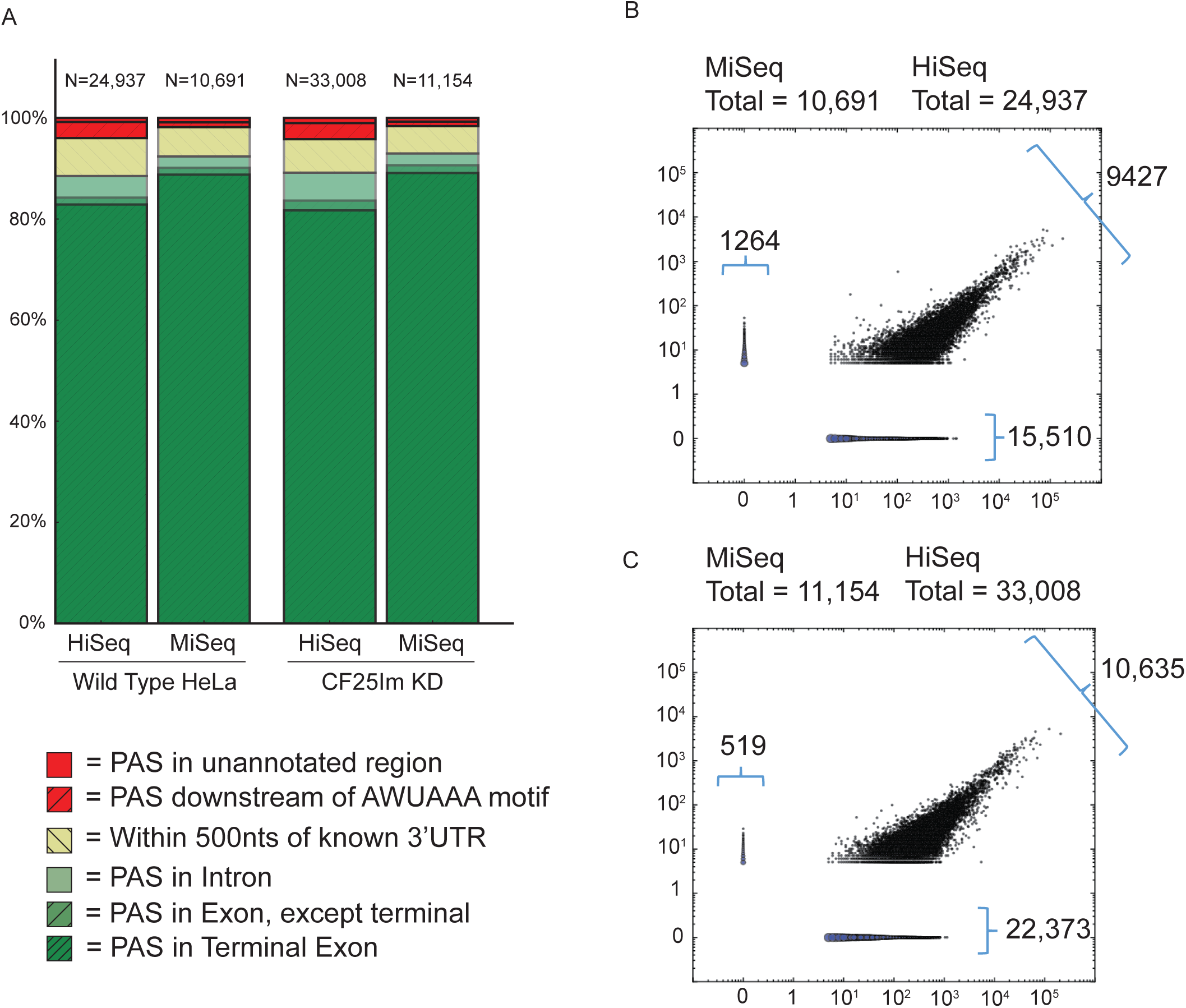
Characterization of Poly(A) sites discovered by Poly(A)ClickSeq. **A)** Bar charts show the overlap of detected poly(A) sites with UCSC knowngene annotations. Poly(A) sties were either found in known exons (Green), less than 500nts upstream of known 3’UTR termini (Yellow), or in unannotated regions (Red). Comparisons between wild-type and CF25Im KD HeLa cell-lines are shown when analysed either using a HiSeq or MiSeq. Scatter-plots comparing the poly(A) site frequencies found by HiSeq and MiSeq analysis for **B)** wild-type and **C)** CF25Im KD HeLa cell-lines. The number of sites in each category are indicated.

When compared to the poly(A) database(54), a total of 20,856 (83.6%) and 26,172 (79.3%) of the detected poly(A) sites for the wild-type and CFIm25 KD datasets respectively mapped over or within 10 nucleotides of the previously annotated sites (**STable 2**). Many of the unannotated poly(A) sites were found to map to mitochondrial genes, highly duplicated loci (e.g. GAGA antigen family) and transposons including LINEs (e.g. Tigger) and SINEs such as *Alu* elements, a large number of which were found within intronic sequences. A large number were also found to be likely uncharacterized pPASs or alternative terminal exons, not currently annotated in the poly(A) database.

#### Replicate sequencing using a MiSeq recapitulates the HiSeq results

PAC-Seq provides an efficient and inexpensive methodology for generating NGS libraries to be sequenced using HiSeq platforms. However, the cost of NGS still remains relatively high and is potentially prohibitive in the analysis of a large number of samples. To determine whether we could obtain the same quality data, but by using a MiSeq platform, we re-sequenced our HeLa cell libraries obtaining 1×250 bp reads. We obtained 880K to 1.51M reads per dataset (**STable 3**), corresponding to 3.5% of the data obtained using the HiSeq. We performed an identical analysis of poly(A) sites (requiring 5 reads to be mapped per poly(A) site, with 5 non-primer-derived A’s, and in at least two replicates) and found a total of 10,691 poly(A) sites in the control-siRNA treated HeLa cells and 11,154 in the CFIm25 KD cells. The distribution of these sites were very similar to that found for the HiSeq data (**Fig. 3A**), with only a small increase in the proportion of poly(A) sites found in terminal exons being observed. This indicates that the less abundant poly(A) sites found only in the high-coverage HiSeq data are enriched for non-canonical poly(A) sites occurs at sites other that the terminal exon.

Calculating the Pearson Correlation coefficient between the HiSeq and MiSeq datasets for the frequencies of mapped reads at each unique poly(A) site returns R values of 0.89 for wild-type HeLa cell and 0.89 for the CFIm25 KD cells. Moreover, as can be seen in the scatter plots in **Fig. 3B** and **Fig. 3C**, the events that were either discovered in the MiSeq but not the HiSeq dataset (and *vice versa*) were the least abundant reads. This demonstrates that we are retaining a large portion of the high-confidence, high-abundance poly(A) sites, despite only mapping 5% the number of processed reads. Therefore, the MiSeq is still sufficient to reproducibly capture the majority of poly(A) events and will therefore be suitable in a broad range of applications, despite the compromise in high-throughput, in a much more cost-effective manner.

#### Poly(A) site choice is promoted by CFIm25 in a UGUA-dependent manner

CFIm25 has previously been implicated in the regulation of the poly(A) cleavage site selection but the mechanism is poorly understood. CFIm25 has been shown to have a preference for UGUA motifs (55) and proximal poly(A) sites have been found to contain elements that do not adhere to consensus as closely as distal poly(A) site motifs do (56). Given that PAC-seq provides an exact polyadenylation site, we decided to explore the relationship of these sequence elements in our datasets. By comparing the control-siRNA treated and CFIm25 knockdown cell-lines, we find a greater number of total poly(A) sites upon CFIm25 KD, despite the fact that we obtained fractionally fewer reads in these datasets (**STable 2**). Moreover, while a slightly higher percentage of poly(A) sites are found in annotated genes (88.5% vs. 89.2%), a slightly smaller percentage of these are found in the terminal exon (82.9% vs. 81.7%). Similarly, a smaller proportion of the poly(A) sites in the CFIm25 KD cells overlap with previously annotated sites in the poly(A) database. Together, these trends may reflect a general role for CFIm25 in specifying the correct PAS (e.g. most consensus) and that a broader range of non-canonical sites become permissive upon CFIm25 knockdown. This hypothesis was explored further.

We first clustered detected poly(A) site in our datasets so that two or more sites found within 10 nts of one another were considered to be same poly(A) site. Next, using the UCSC *knowngene* annotations (53), we considered only poly(A) sites that were found in the terminal exons. For the HeLa cells, from a total of 9841 individual mRNAs, we found 3388 mRNAs with two or more poly(A) sites containing a total of 7651 unique poly(A) sites (**Fig. 4A**). 1776 mRNAs were determined to exhibit significant (greater than 20% change) APA upon CFIm25 KD. Of these mRNAs, 1430 exhibited 3´ UTR shortening (**Fig. 4C**) and 346 exhibited 3’ UTR lengthening (**Fig. 4E**). The large number of shortened genes is highly consistent with previous studies by us and others.

**Figure 4:**
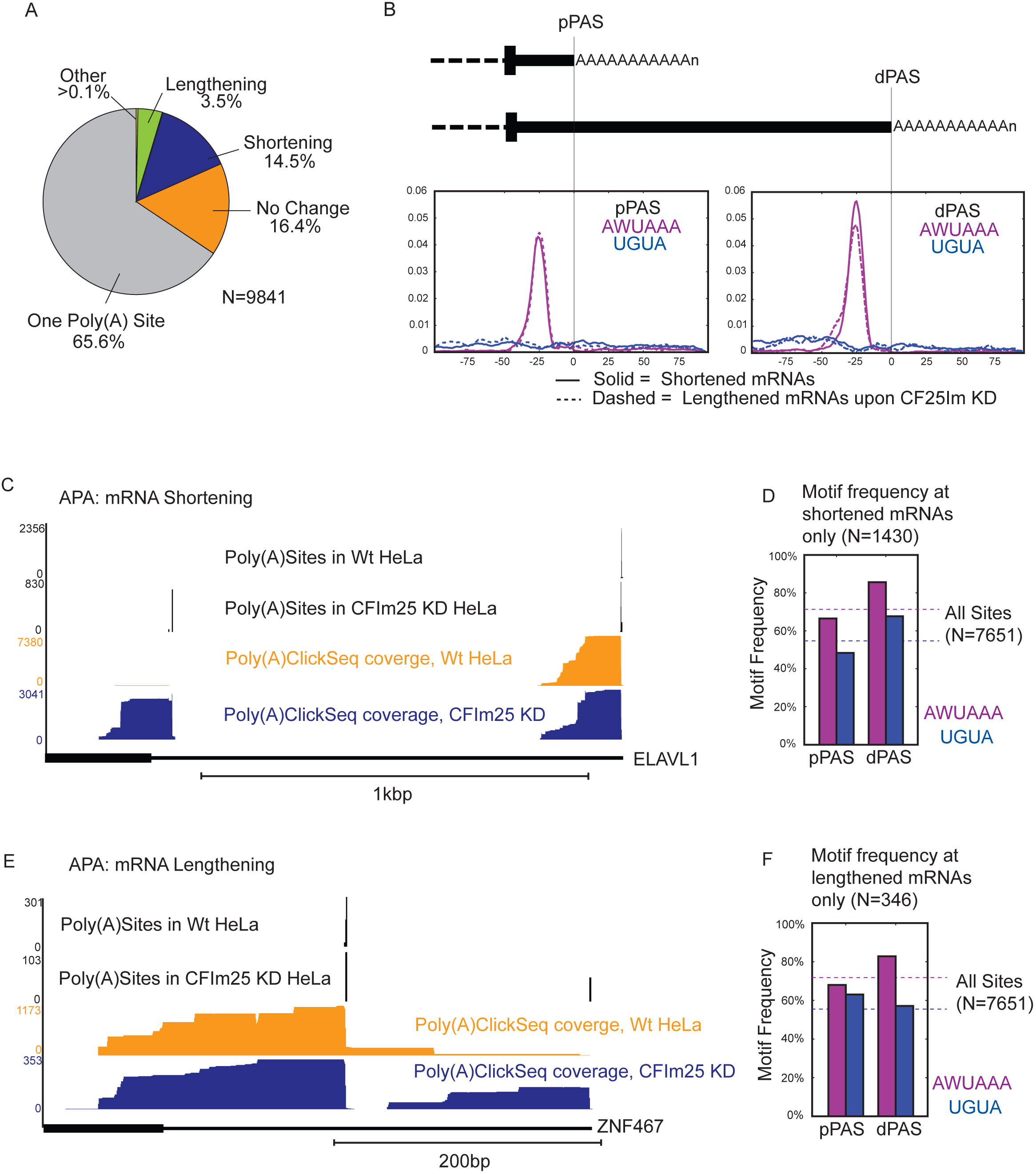
Alternative poly-adenylation upon CF25Im knock-down. The poly(A) sites found in wild-type and CF25Im KD HeLa cells lines were compared and revealed whether mRNAs had one or multiple poly(A) sites within annotated terminal exons, and whether CF25Im knock-down resulted in the lengthening of shortening mRNA transcripts. Frequency of poly(A) sites in these categories are shown in **A)**. **B)** Motif enrichment analysis using the *MEME* suite revealed that AWUAAA and UGUA motifs were enriched upstream of both proximal (pPAS) and distal (dPAS) poly(A) sites. *Centrimo* analysis showed that AWUAAA was found 20-40 nucleotides upstream of PASs while UGUA was dispersed. **C)** Poly(A)Site and coverage tracks are shown for an example of a gene (ELAVL1, Human Antigen R) exhibiting 3’UTR shortening upon CF25Im KD. **D)** The frequency of AWUAAA and UGUA motifs found within 100nt upstream of both proximal (pPAS) and distal (dPAS) poly(A) sites only for shortened mRNAs are shown. **E)** Poly(A)Site and coverage tracks are shown for an example of a gene (ZNF467, Zinc Finger Protein 467) exhibiting 3’UTR lengthening upon CF25Im KD. **F)** The frequency of AWUAAA and UGUA motifs found within 100nt upstream of either both proximal (pPAS) and distal (dPAS) poly(A) sites only for lengthened mRNAs are shown.

The differential usage of 3´ poly(A) cleavage sites is poly-factorial, but has been demonstrated to be promoted by the presence of at least two PAS motifs: AWUAAA and UGUA (4). *DREME* analysis(47) of these sites confirmed that these motifs were significantly enriched in the regions upstream of the detected poly(A) sites. To determine whether the choice of poly(A) cleavage site was altered by CFIm25 in a manner dependent upon these motifs, we quantified the number of poly(A) sites containing AWUAAA and UGUA motifs <100 nts upstream. For all 7651 sites, we found that 71.9% and 56.0% contained AWUAAA and motif UGUA motifs respectively. Using *CentriMo*(*48*), we found that the AWUAAA motifs are strongly enriched between 20 and 40 nts preceding the PAS, but that UGUA motifs show little positional preference (**Fig. 4B**). This was true for both proximal and distal sites, regardless of whether CFIm25 KD induced lengthening or shortening of 3´UTRs.

To investigate why most mRNAs exhibited 3´UTR shortening while a small group of others presented lengthening in response to CFIm25 knock down, we analyzed the frequency of the AWUAAA and UGUA motifs found upstream of both the proximal (pPAS) and distal (dPAS) poly(A) sites for both lengthened and shortened mRNAs. We find that distal sites are relatively enriched for AWUAAA motifs (>80%) regardless of whether CFIm25 KD induced lengthened or shortened 3´UTRs (**Fig. 4B, D and F**). Interestingly however, we observe a different trend for UGUA whereby UGUA motifs are enriched in the distal sites of mRNAs that are shortened after CFIm25 KD, but enriched in the proximal sites of mRNAs are lengthened after CFIm25 KD. This implies that selection of these poly(A) sites are promoted by CFIm25 in a UGUA-dependent manner and that this enhancement was lost upon CFIm25 KD resulting in an increase in the expression of less optimal alternative poly(A) sites. If the UGUA site is more prevalent in the distal position, then knockdown of CFIm25 reduces the utilization of this site and so the mRNAs are shortened. Conversely, if the UGUA site was found in the proximal position then that mRNA is lengthened. These trends were conserved when only considering mRNAs with only 2 PASs and at >20%, >50% and >80% APA strength (**SFig 5**). Collectively, these results exemplify how PAC-seq datasets can be leveraged using motif analysis tools and provide additional insight into how CFIm25 is regulating poly(A) site choice.

#### Poly(A)-ClickSeq analysis of Drosophila S2 cells

We sought to further determine to what extent CFIm25 regulation of alternative polyadenylation is conserved in invertebrate species and how effective PAC-seq is in the analysis of a novel RNA dataset. Using dsRNA targeting the Drosophila orthologue of CFIm25 (CG3689), we knocked down CFIm25 in S2 cells to a degree exceeding 90% (**SFig 4B**). This level of knockdown in mammalian cells is sufficient to trigger genome-wide 3´UTR shortening in human cells and, in our experience, is sufficient to mediate a spectrum of loss of function phenotypes in S2 cells (57-61). Total cellular RNA from both mock-treated cells and CFIm25 KD cells was harvested in triplicate and used to generate PAC-Seq libraries using the same procedures as those described above. Libraries were sequenced on a MiSeq obtaining a total of 2.39M and 3.09M total processed reads, on average, for mock and CFIm25 KD S2 cells. Using the same mapping parameters as above (requiring 5 reads with poly(A) tails 25 nts of longer to be found in at least two of the three replicates) we found 6910 poly(A) sites in the mock cells and 7473 in CFIm25 KD cells (**STable 4**). Our dataset compares well with the poly(A) sites determined in a previous analysis (62), with 62.0% and 53.8% of our poly(A) sites found within 10nts of the previous data. Similar to HeLa cells, these poly(A) sites are primarily found at the terminal exons of annotated genes (79.1% and 79.2%) (**STable 4 and Fig. 5A**). However, in each dataset a proportionally larger amount (16.2% and 15.7% in Wt and CFIm25 KD) were found to map within 500nts downstream of the annotated genes in the UCSC database suggesting the S2 cells express a significant amount of mRNA that are longer than annotated or that the annotations of the 3’ ends of genes are imprecise. A small proportion of the poly(A) sites were found in unannotated regions, 146 (1.9%) and 174 (2.2%). Again, 54 and 67 of these contain AWUAAA motifs within 100nts upstream of the poly(A) sites reported here, indicating that a significant proportion of these corresponded to likely bona fide poly(A) sites.

**Figure 5:**
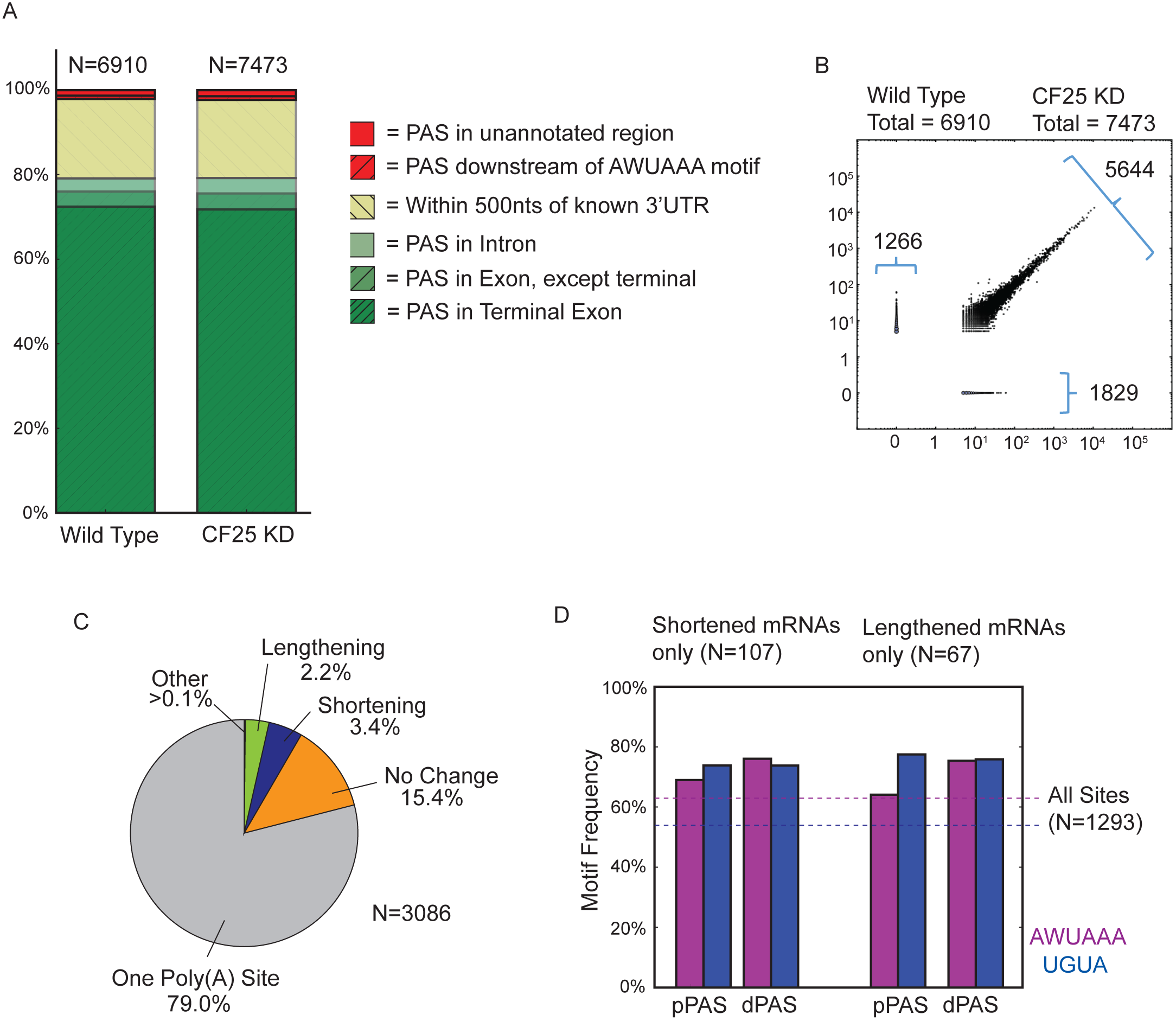
Poly(A)ClickSeq analysis of Drosophila melanogaster cells in culture reveal litte effect of CF25d known-done upon poly(A) site selection. **A)** Bar charts show the overlap of detected poly(A) sites with UCSC knowngene annotations. Poly(A) sties were either found in known exons (Green), less than 500nts upstream of known 3’UTR termini (Yellow), or in unannotated regions (Red). **B)** A scatter-plot comparing the frequency of poly(A) sites in both wild-type and CF25d KD S2 cell is shown revealing very high correlation between the two datasets (R=0.99). **C)** The effect of CF25d knock-down upon poly(A) site diversity is limited in Drosophila. Frequency of poly(A) sites in the same categories as described for **Fig 4B** are shown. **D)** The frequency of AWUAAA and UGUA motifs found within 100nt upstream of both proximal (pPAS) and distal (dPAS) poly(A) sites are shown for both shortened and lengthened mRNA.

We next characterized any changes in poly(A) site selection upon CFIm25 KD. Unlike the observation of broad APA in human cells in response to CFIm25 knockdown, we observed less changes in poly(A) site position and frequency when the fly orthologue is knocked down (**Fig. 5B**, Pearson coefficient = 0.99). Moreover, a much smaller proportion of genes contained multiple poly(A) sites (648 out 3086 genes) than was found for the HeLa cells (**Fig. 5C**). From these, a small percentage of these exhibited significant APA upon CFIm25 KD (greater than 20% change in PAS abundance - 107 shorted, 67 longer, 5 both).

Nonetheless, for the few genes that did exhibit APA, we analyzed enrichment of the AWUAAA and UGUA motifs. Both these motifs were found to be significantly enriched upstream of the poly(A) sites (72.0% and 76.5%) and their positional distribution was found to be similar to that observed for HeLa cells (**SFig. 6**). However, while we observe a similar enrichment for AWUAAA motifs in the distal sites, we now also observe an enrichment for UGUA (**Fig. 5D**). Importantly however, this enrichment is the same regardless of whether mRNAs are lengthened or shortened upon CFIm25 KD (in contrast to the HeLa cell data, compare to **Fig. 4D** and **F**). This implies that APA in S2 cells is less dependent upon CFIm25 for poly(A) site choice.

### DISCUSSION

As the applications of next-generation sequencing grow and diversify, a key challenge will be developing cost-effective, robust, and sensitive methods for the generation of targeted cDNA libraries. Here we presented a simple, quick and inexpensive method for the generation of next-generation sequencing libraries called Poly(A)-ClickSeq (or PAC-seq) that specifically enriches for the junction of the 3´ UTR and poly(A) tail junction. We demonstrated that we could recapitulate the findings of previous analyses of the poly(A) landscape in both human and *Drosophila* cell-lines. As well as confirming the presence of previously annotated transcripts termination sites, PAC-seq was also able to identify novel poly(A) sites that are likely *bona fide* given their proximity to AWUAAA.

Using this approach, we also demonstrate that poly(A) sites that are down-regulated upon CFIm25 knock-down are relatively enriched for the UGUA motif. While the majority of these downregulated sites are at the distal poly(A) site resulting in 3’UTR shortening, there was a small group of transcripts that underwent 3’UTR lengthening, which correlated with the enriched UGUA motif is located at the proximal poly(A) site. These two observations generate a simplified model where reduced expression of CFIm25 will result in loss of enhancement of poly(A) sites that are rich in UGUA causing the usage of other poly(A) sites within a given transcript. This model is simpler in that it does not require that CFIm25 functions as a repressor of poly(A) site selection but rather is always an enhancer of cleavage and polyadenylation, which is consistent with its originally postulated function as an essential CPA member.

Our method provides a number of advantages over other popular approaches. The first is that no sample preparation or purification is required. We demonstrated here that poly(A) sites can be sequenced directly from total cellular RNA extracts without enrichment for polyadenylated RNAs or removal of ribosomal RNAs (for example). This has three important consequences: i) these enrichment/depletion steps are time-consuming and their cost can be significant; ii) enrichment/depletion steps can potentially impart significant bias leading to uneven sequence coverage, and can inadvertently obscure potentially interesting species (such as rRNA degradation products); and iii) library generation is markedly simplified, reducing manipulation and loss of precious samples. Together these factors may allow for the use of PAC-seq in highly challenging biological contexts such as the poly(A) profiling directly from tumor biopsies.

A second key advantage is that, similar to ClickSeq, PAC-seq does not require RNA sample fragmentation. There are few available methodologies that remove the fragmentation steps of NGS library synthesis. Removing this step again simplifies sample preparation, and also avoids the biases that can arise due to RNA fragmentation protocols and subsequent adaptor ligation. This advantage also removes any need for specialized equipment beyond standard laboratory items. Another advantage is that we use non-anchored poly(T) primers, allowing non-primer-derived As to be found in the final RNAseq reads. As described in the *methods* section, this allows for an additional quality filtering protocol that substantially improves confidence in reported poly(A) tails. Moreover, the distributions of poly(A) lengths can be inferred for each detected poly(A) sites. Poly(A) tail length is an important variable affecting RNA stability and half-life. Therefore, PAC-seq may also be used to assess site-specific changes in poly(A) tail lengths.

Although we did not explore this possibility in this manuscript, the click-ligated adaptors can also be designed to contain single-molecule indexes, similar to the PrimerID strategies used to sequence HIV protease(63). This can allow for sequence error correction and perhaps more importantly, for assessment of PCR mediated duplication bias. For some samples, it may be necessary to perform many rounds of PCR amplification in order to generate enough substrate to load onto an Illumina flowcell. By including single-molecule indexes in the click-adaptor, over-sampling errors can be corrected.

Overall, PAC-seq is a simple, quick and cheap method for NGS library generation that captures the 3´UTR/poly(A) tail junction with high efficiency resulting in a reduced need for sequence depth. From our initial HiSeq dataset, approximately 50% of the total raw sequences reads were utilized to the final analysis. While saving on cost, this also allows for a single experiment with multiple replicates to be performed on a single MiSeq flowcell. The current v3 MiSeq kit can yield ~25 million read under optimal conditions. This would allow over ten replicates of a single experiment at a coverage of 2 million reads per dataset. This coverage depth is sufficient for analyzing even highly complex genomes such as in human cells.

## ACKNOWLEDGEMENTS

We thank Thomas Wood, Steve Widen and Jill Thomson from the UTMB Next-Generation Sequencing core and Gillian Lynch from the SCSB computing facility for support. We are indebted to Todd Albrecht, Mariano Garcia-Blanco, Shelton Bradrick and members of their lab for sharing lab space, helpful discussions and other research support.

